# Vascular diversity in Fabaceae: evolutionary and ecological insights from a globally distributed lineage

**DOI:** 10.64898/2026.03.20.713002

**Authors:** John Karlo C. Saddoy, Israel L. Cunha-Neto

## Abstract

The vascular system is central to plant ecology and evolution. Here, we show that more than 100 species across 27 genera and four subfamilies of Fabaceae have evolved atypical vascular architectures and that these species occur in all biogeographical regions except Antarctica. Because Fabaceae includes many ecologically and economically important species exhibiting these novel vasculatures, the family emerges as an ideal system for assessing the implications of vascular innovation in both fundamental and applied research.

## 1. Introduction

Fabaceae species exhibit diverse growth forms, including trees, shrubs, climbers, herbs, and hydrophytes (Siddiqui et al. 2025). Specifically, the family is the third-largest lineage of climbers (Sperotto et al. 2023), ranging from delicate herbaceous vines to some of the largest lianas (woody vines) thriving in temperate and tropical forests (Acevedo-Rodríguez 2015 onwards). These climbing plants, like other gymnosperm and angiosperm climbers, are known for their specialized vascular system that enables them to climb without snapping (Carlquist 1991; Angyalossy et al., 2012, 2015). Specialized vascular traits in these vines include the co-occurrence of narrow and wide water-conducting cells, abundant soft tissues intertwined with rigid tissue, and, at the most extreme, the appearance of “vascular variants”—which are diverse vascular architectures that deviate from the solid cylinder of wood and inner bark observed in most woody plants (Cunha Neto 2023).

The combination of vascular traits forming the adaptive liana vasculature has been termed “lianescent vascular syndrome” (Angyalossy et al. 2012; 2015), a concept further validated through quantitative comparative analysis of wood and bark traits from Fabaceae trees and lianas (Leme et al. 2021). In addition, several studies examining the presence of vascular variants in stems and roots have enhanced our understanding of vascular diversity in Fabaceae (Kraus & Basconsuelo 2009; Nejapa et al. 2022; Rajput et al. 2023; Rami et al. 2025) and in seed plants altogether. In particular, studies of vascular biology in Fabaceae have indicated that vascular variants occur both in lianas and self-supporting species (e.g., *Dalbergia* trees; Nair and Mohan Ram 1990), highlighting functions beyond those associated with climbing plants (e.g., stem flexibility). However, we still don’t fully understand how and why these novel vasculatures evolved because studies integrating ecology, evolution, and developmental biology (eco-evo-devo) within lineages with vascular variants remain scarce. The distribution of vascular variants across species with diverse growth forms and geographical ranges makes Fabaceae an excellent group for filling this gap.

Among the nine patterns of vascular variants reported in Fabaceae (Cunha Neto 2023), ectopic cambia (EC)—the repeated formation of vascular increments of water-conducting tissue (secondary xylem) and sugar-conducting tissue (secondary phloem) on a single stem or root—is by far the most ecologically and economically significant. Ectopic cambia result from the natural occurrence of multiple cambia (each producing a vascular increment) as opposed to the single cambium seen in the “typical growth” of most woody plants, which produces one continuous cylinder of secondary xylem (wood) and secondary phloem (inner bark). Ectopic cambia characterize the woody stems and roots of trees, shrubs, and lianas in the family Fabaceae, as well as 44 other families of gymnosperms and angiosperms (Cunha-Neto 2026). In Fabaceae, EC occur in essential crops (lima bean), ornamentals (wisteria, Jade vine), and noxious weeds (kudzu) (Obaton 1960; Yang et al. 2022). Some of them are endemic to specific regions (e.g., Jade vine in the Philippines), others are distributed across multiple continents (e.g., *Dalbergia*), or found across temperate and tropical regions (e.g., *Mucuna*), indicating their diverse biogeographical and ecological distributions. However, a comprehensive assessment of the diversity, evolution, and biogeographical distribution of species with EC is lacking in Fabaceae, despite the existence of a family-level phylogeny (Azani et al. 2017) and morphological data indicating their occurrence for over a century (De Bary 1877; Solereder et al. 1908). Interestingly, the presence of EC does not necessarily require developmental studies, as this phenomenon produces macromorphologies that can be identified even in herbarium/wood collection specimens, including images in online databases.

Here, we begin exploring the ecology and evolution of vascular diversity in Fabaceae, presenting the first assessment of EC evolution within the family, and offering insights into the evolutionary transitions of vascular variants across plants with different habits and geographic distributions. This research comes in part from a new initiative that established permanent plots in the Mount Makiling Forest Reserve (MMFR), Los Baños, Philippines, to support long-term research into the ecology and evolution of climbing plants at a local scale. Furthermore, we introduce the “Plant Vascular Variants Database,” a curated, continuously updated resource for the study of vascular variants (Cunha Neto 2026).

## 2. Methods

To build our database of ectopic cambia (EC) occurrences across Fabaceae and closely related specimens, we used morpho-anatomical images of the cross-sectional area of stems and roots from species obtained from field-collected samples and from the literature, including online botanical databases (Supplementary Table 1). For newly collected specimens from the permanent plots in the MMFR, Los Baños, Philippines, we employed standardized procedures in vascular biology (Angyalossy et al. 2016; Barbosa et al. 2021) to generate macroscopic images and characterize EC development. The database for EC distribution across Fabaceae is available in Supplementary Table 1 and will be continuously updated as an independent module (Saddoy and Cunha-Neto, 2026) within the “Plant Vascular Variants Database” (Cunha-Neto 2026).

We examined the evolutionary history of EC and growth forms using phylogenetic comparative methods. To that aim, we integrated our anatomical dataset (Supplementary Table 1) with a Fabaceae phylogeny modified from Azani et al. (2017). For ancestral state reconstructions (ASR), we performed two complementary analyses. First, a “species-level ASR” was performed by matching species names in the anatomical dataset directly to tip labels in the phylogeny for which exact matches were available. Second, a “genus-level ASR” was conducted, including genera represented in the anatomical dataset but absent as the same species in the phylogeny; by assigning EC to congeneric species present in the phylogeny, we maximized generic coverage by five genera (Supplementary Table 2). Ancestral states and independent origins of EC were quantified from the “genus-level ASR” using stochastic character mapping in *phytools* (Revell 2012). We used Pagel’s test (1994), as implemented in *phytools* to test for correlated evolution between EC and the liana growth form. We selected representative species to plot on the map of floristic biogeographical realms (Liu et al., 2023) to visualize the distribution of species with EC across the globe.

## 3. Results

We found ectopic cambia (EC) in 109 species of Fabaceae, representing 27 genera and four subfamilies, which occur in all biogeographical realms, except Antarctica (Fig. 1; Supplementary Table 1). The Neotropical realm has the highest number of species with EC (46) and is followed by the Indo-Malesian (24), Australian (19), Holarctic (10), Afrotropical (5), and Palaeotropical (5) realms. The Indo-Malesian realm has the highest number of endemic genera with EC (7), followed by the Neotropical (6), Australian (3), Holarctic (2), and Afrotropical (2) realms.

**Figure 1.**
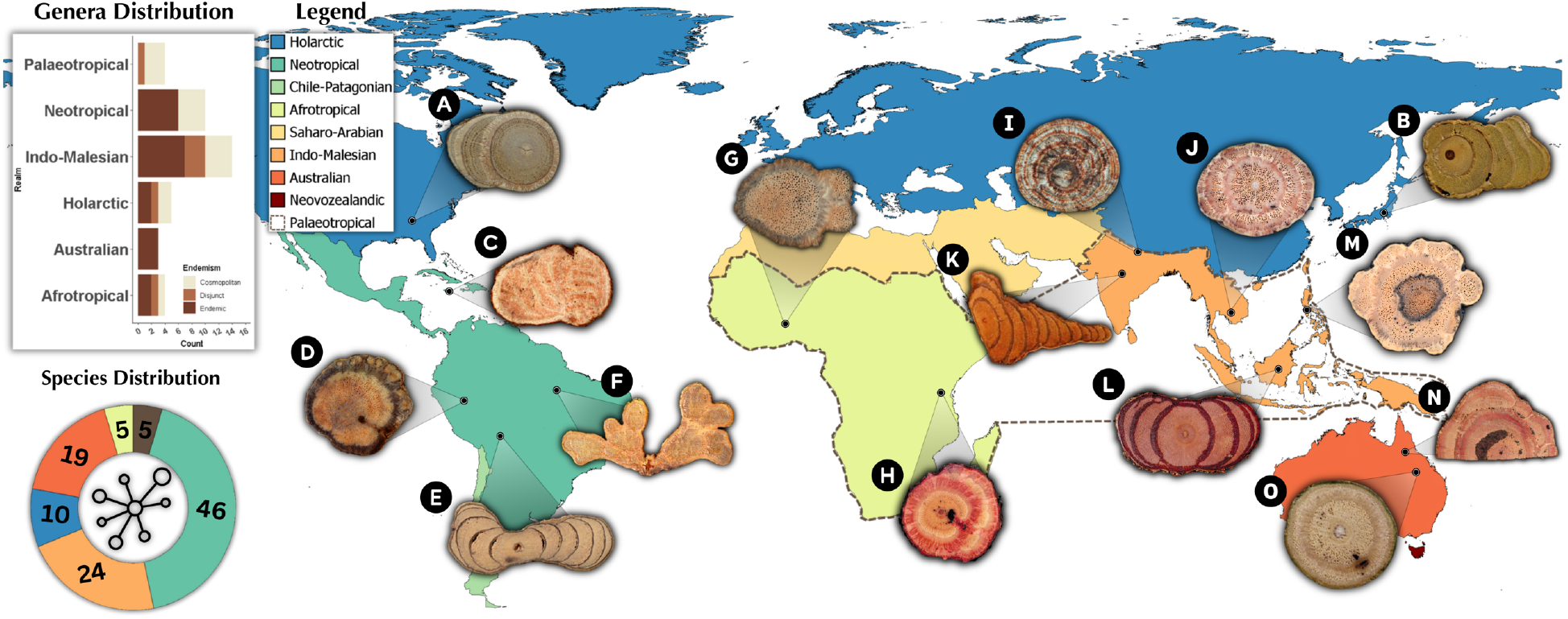
Biogeographical distribution of Fabaceae species with ectopic cambia (EC). Images illustrate stem cross-sections. Stem diameters range from 2-10 cm. (A) *Wisteria sinensis* (Nejapa et al. 2022; Crown 2021). (B) *Wisteria floribunda* (Nejapa et al. 2022; Crown 2021). (C) *Entada gigas* (reproduced with permission from Acevedo-Rodríguez 2015 onwards). (D) *Phaseolus lunatus* (Rajput et al. 2023; reproduced with permission from Elsevier). (E) *Machaerium latifolium* (reproduced with permission from Acevedo-Rodríguez 2015 onwards). (F) *Schnella microstachya* (reproduced with permission from Acevedo-Rodríguez 2015 onwards). (G) *Lablab purpureus* (Yang et al. 2022; CC BY 4.0). (H) *Derris trifoliata* (reproduced with permission from Zich et al. 2020; CANBR 1989). (I) *Mucuna macrocarpa* (Yang et al. 2016; reproduced with permission from *TAIWANIA*). (J) *Pueraria montana* var. *lobata* (John Karlo C. Saddoy). (K) *Spatholobus suberectus* (Qin et al. 2020; CC BY 4.0). (L) *Dalbergia ferruginea* (reproduced with permission from Zich et al. 2020; CANBR 1980). (M) *Strongylodon macrobotrys* (John Karlo C. Saddoy). (N) *Austrosteenisia stipularis* (reproduced with permission Zich et al. 2020; CANBR 1989). (O) *Brachypterum koolgibberah* (reproduced with permission from Zich et al. 2020; CANBR 1995). Note: The Palaeotropical realm (sometimes referred to as the Old World tropics) encompasses both the Afrotropical and Indo-Malesian realms. Generic endemism in biogeographical distribution is based on Huang et al. (2025), in which endemic genera are clustered within one realm, disjunct genera are separated by wide or significant geographical boundaries, and cosmopolitan genera are found across multiple continuous realms.

Ectopic cambia is disproportionately more abundant in lianas with a record of 87 species, but was also recorded in 17 shrubs (e.g., *Daviesia* and *Sophora*), four trees (*Dalbergia, Koompassia, Machaerium*), and one herbaceous vine (*Pachyrhizus*) (Supplementary Table 1). Although EC is commonly reported in stems, 17 of the 109 species also exhibit EC in their roots (including the storage roots of kudzu); however, information on root vasculature is mostly unknown. *Machaerium* is the genus with the highest number of species with EC recorded to date, while eight genera have only one species recorded (Supplementary Table 1).

The ancestor of Fabaceae was reconstructed as a tree with typical growth (Fig. 2A). In the “genus-level ASR” (Fig. 2A), EC evolved three times in the outgroup (Polygalaceae) and several times independently in Fabaceae: once in Cercidoideae, Dialioideae, and Caesalpinioideae, and more than 10 times in Papilionoideae, with several reversals to typical growth. Pagel’s test of correlated evolution showed that the evolution of EC depends on the evolution of the liana habit in Fabaceae (p < 0.005; see Supplementary Figure 1 for a mirrored tree showing stochastic mapping of vascular patterns and growth forms).

**Figure 2.**
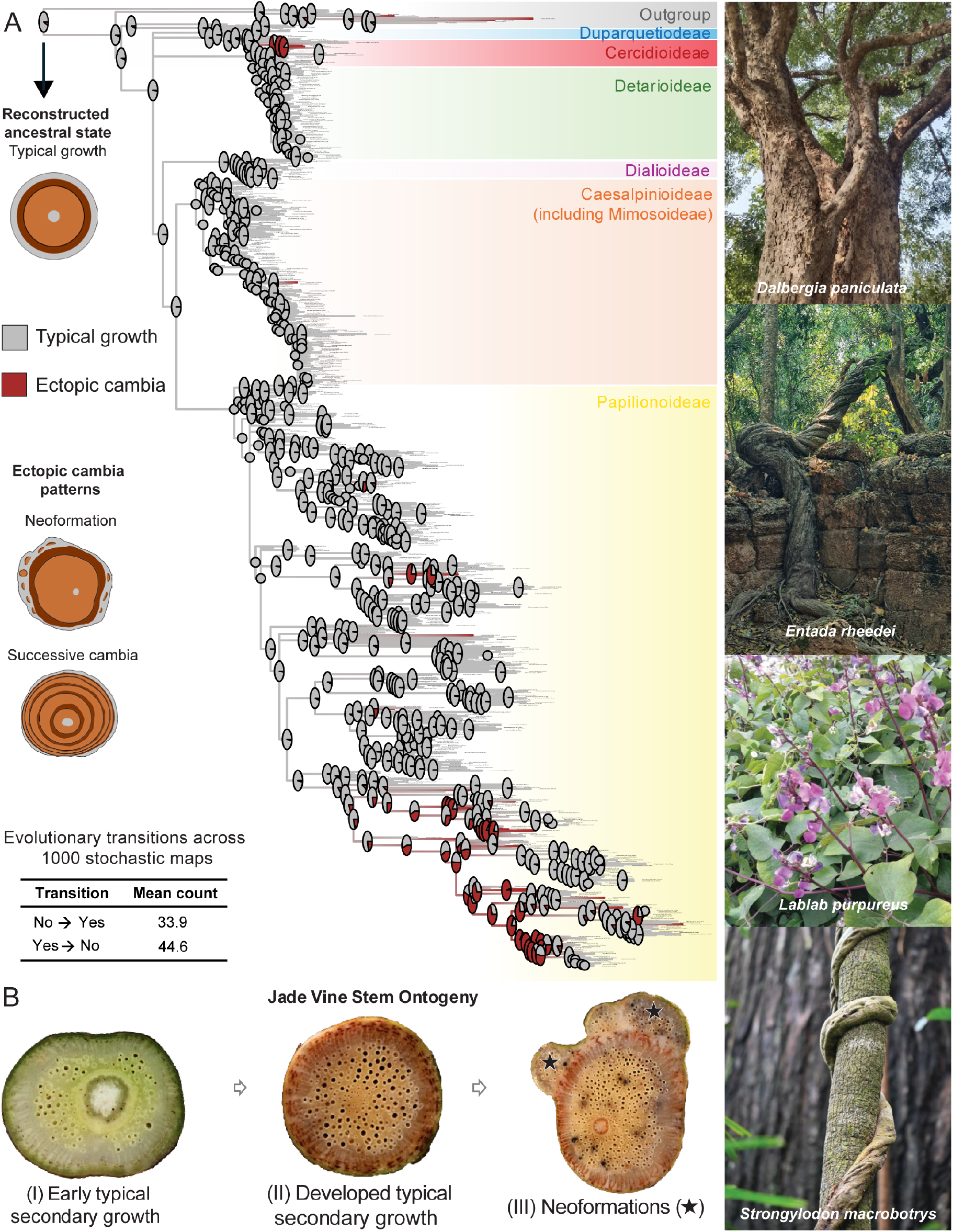
Evolution and anatomical diversity of ectopic cambia (EC) in Fabaceae. (A) Stochastic character mapping of EC using the genus-level dataset, showing multiple independent origins of EC from ancestors with typical growth. Ectopic cambia is defined as a binary character, lumping different EC morphotypes (e.g., successive cambia, neoformation). In the drawings, tan indicates the secondary xylem (wood); brown indicates the secondary phloem (inner bark), and the vascular cambium is represented by the black lines between these two tissues. Ectopic cambia were found in four subfamilies as different morphotypes (neoformations and successive cambia) that evolved in trees, shrubs, and lianas (see images on the right). The number of independent origins is summarized from 1000 stochastic maps. (B) The Jade vine *(Strongylodon macrobotrys*) contains EC in roots and stems; stem development shows typical secondary growth (I), continues in mature stems, from which EC may appear at any point in time, arising from cortical parenchyma. Stem diameter: I = 8.7 mm, II = 15.7 mm, and III = 32.3 mm. Image credit: *Entada rheedei* (iNaturalist, photo by Heidi Clark; CC BY-NC 4.0), *Dalbergia paniculata* (iNaturalist, photo by Sonu Kumar; CC BY-NC 4.0), *Lablab purpureus* (iNaturalist, photo by along2022; CC BY-NC 4.0).

We characterized the developmental anatomy of EC in the charismatic Jade vine (*Strongylodon macrobotrys*) for the first time. Jade vine is a Philippine endemic liana and popularly cultivated in botanical gardens worldwide. We found EC in the roots and stems of all four specimens collected at the MMFR permanent plots (Supplementary Table 3). Ectopic cambia developed during late secondary growth, after a significant amount of typical wood and inner bark had been produced (Fig. 2B). Ectopic cambia form in the cortex and are arranged discontinuously, creating isolated vascular units (neoformations) (Supplementary Figure 2).

Ectopic cambia development produces different macroanatomical patterns in Fabaceae (Fig. 2A). Out of the 109 species, 78 have concentric and continuous vascular increments in the form of “successive cambia” rings, while the rest display discontinuous or isolated vascular units known as “neoformations” (Supplementary Table 1). Ectopic cambia in some taxa are developmentally variable and will require further studies to disentangle their developmental diversity (e.g., *Entada*; Supplementary Figure 3).

## 4. Discussion

Our findings support ectopic cambia (EC) as a convergent trait in seed plants (Cunha-Neto and Onyenedum 2023) and, for the first time, systematically demonstrate their wide distribution across biogeographical realms. The distribution of lianas with EC across distinct geographical regions corroborates the presence of vascular variants as a high adaptive strategy, which may be related to increased stem flexibility, higher hydraulic capacities, mechanical stiffness (Rowe & Speck 2005), and injury repair (Fisher & Ewers 1989). Nevertheless, the observation of constitutive EC in diverse self-supporting plants (trees, shrubs, and herbs) suggests distinct adaptive advantages, which may include physiological responses linked to microenvironmental factors (e.g., mangrove trees; Robert et al., 2011) or alternative hydraulic pathways (e.g., Caryophyllales; Cunha-Neto et al., 2024). Induced/regenerated cambia have also been observed in lianas and trees (e.g., girdled poplar trees; Zhang et al., 2011), particularly in Fabaceae, where previous research has shown they are triggered by physical barriers (Rajput 2003), herbicide treatment (soybean; Struckmeyer et al. 1976), and fungal diseases (Li et al. 2018). These observations demonstrate that constitutive EC, which may share similar developmental mechanisms with regenerated cambia (Cunha Neto et al., 2026), is a crucial vascular innovation in the evolutionary history of woody plants and a regeneration mechanism that may be vital for plant adaptation.

Previous studies on evolution of development of vascular variants (including EC) showed that, in several cases, evolutionary transitions from typical growth to vascular variants have occurred in lineages containing lianas or descending from a liana ancestor in four of the five families studied to date (i.e., Bignoniaceae, Malpighiaceae, Malvacee, and Sapindaceae) (Pace et al., 2009; Luna-Márques et al., 2021; Quintanar-Castillo et al. 2022, 2025; Cunha Neto et al. 2025), while in one case (i.e., Nyctaginaceae), they are widespread, regardless of plant habit (Cunha Neto et al., 2022). Fabaceae is the sixth family investigated using this approach, showing that transitions from typical growth to EC are not only more common in lianas but also correlated with their evolution, although many trees, shrubs, and herbs also exhibit these alternative forms. The repeated transitions from typical growth to EC in Fabaceae may be linked to its status as the third-largest angiosperm family and to the ease of genetic changes that enable these evolutionary transitions. The first study exploring the genes involved in EC formation was conducted in Fabaceae species, the Japanese wisteria, and the results suggest that EC development may require changes in conserved genes linked to typical vascular development (e.g., *KNOX*) (Cunha Neto et al., 2026). However, much remains to be elucidated about the gene regulatory networks underlying plants’ ability to create EC (naturally or induced).

To advance our knowledge of fundamental and applied plant vascular biology, we identify two areas for continuous collaboration. First, we acknowledge the critical role of herbaria, wood collections, and online repositories as essential resources not only for traditional investigations in plant systematics and conservation but also for exploring morphoanatomical traits that support ecological and evolutionary research. For example, many species with EC assigned in the present study originated from online databases, including 17 species from the curated works of Dr. Pedro Acevedo-Rodríguez at the Smithsonian Institution (Acevedo-Rodríguez 2015 onwards), and five species from the Australian flora database of Zich et al. (2020). We urge scientists and stakeholders to ensure the continued maintenance and improvement of these resources. In this context, we have also launched the “Plant Vascular Variants Database” (Cunha Neto 2026; https://www.plantgrowthlab.com/plant-vascular-variants-database), a continuously updated repository that centralizes information on vascular variants previously scattered across the literature.

Secondly, we emphasize long-term research in permanent plots as a valuable approach to further exploring plant vascular diversity, particularly in lianas. Inspired by the Liana Project in Panama, including its recent initiative on “The Search for Champion Lianas” (Schnitzer et al., 2025), as well as other pioneering community-level studies on the functional roles of vascular variants (Caballé 1986; Yang et al. 2022; Kaçamak et al. 2025), this investigation represents one of several efforts underway in the newly established plots at the Mt. Makiling Forest Reserve (Los Baños, Philippines), where we are systematically documenting liana ecology in the country for the first time. We encourage similar initiatives in other tropical regions with high liana diversity and expertise in liana biology, such as Brazil, India, and Mexico, as well as in subtropical areas like South Florida, United States, where pioneering research in liana biology has been conducted (Fisher & Ewers 1989).

Overall, our findings support the adaptive role of vascular variants in lianas and highlight the Fabaceae as an excellent group for exploring the importance of alternative vascular forms in plants with diverse growth habits, as well as their relevance for research efforts that connect vascular biology with applied research in plant sciences.

## Author Contributions

I.L.C.N. conceived the original project. J.K.C.S. performed fieldwork. All authors proposed methods, collected and curated data, wrote the original draft, and edited the final version.

## Acknowledgements

We thank Dr. Lerma SJ. Maldia, Director of the Makiling Center for Mountain Ecosystems, CFNR, UPLB, for granting permission to collect Jade vine specimens in Mt. Makiling Forest Reserve (Permit to Conduct Activity, No. FWD-2024 – 121; this permit has an equivalent implementing power to the Gratuitous Permit of the Department of Environment and Natural Resources (Republic Act No. 9147 or Wildlife Resources Conservation and Protection Act). We thank the staff at the LBC Herbarium and Wood Collection (University of the Philippines at Los Baños) for the curation of our Jade Vine samples. We also thank the collaborators and repositories that contributed images. Grammarly (Grammarly, Inc.) was used to check for grammatical errors and improve the clarity of the text.

## Data Availability Statement

The data that support the findings of this study are openly available in the Figshare Repository: 10.6084/m9.figshare.31816603

## Supporting Information

**Supplementary Table 1.** Diversity of ectopic cambia in Fabaceae across different growth forms, biogeographical locations, and plant organs (stem or roots).

**Supplementary Table 2.** Alternative species used to represent the genus for the genus-level ASR analysis in the phylogenetic tree.

**Supplementary Table 3.** Permanent plot-based location and the corresponding morphometrics of collected Jade vines.

**Supplementary Figure 1.** Evolution of vascular architectures and growth forms using the species-level Fabaceae dataset. Ancestral states on the branches are summarized from 1000 stochastic maps, with observed states indicated at the tips. Pagel’s test of correlated evolution showed that ectopic cambia and the liana habit evolved in a correlated fashion.

**Supplementary Figure 2.** Ectopic cambia development in Jade Vine (*Strongylodon macrobotrys* A. Gray). All images are cross-sections. (A-C) Transition from primary to secondary growth. The vascular cambium (ca) connects the vascular bundles, whose positions can still be seen based on protoxylem poles (arrows), in the periphery of the pith (pi). A fibrous pericyclic ring (pf) and secretory cells (with dark content) are observed. (D-H) Mature stem showing secondary growth derived from the original cambium, which produces secondary xylem (sx) and secondary phloem (ph). Note the first increment of ectopic cambia (dashed boxes). Ectopic cambia arise from cortical parenchyma, which can be confirmed based on the position of pericyclic fibers (arrowhead). Note a neoformation with perpendicular cambium (asterisk) orientation in H. Bars: 200 μm (A, F), 100 μm (B, C), 500 μm (D, E), and 50 μm (G, H).

**Supplementary Figure 3.** Unique successive cambia developmental conformation of assessed *Entada* spp. (A) *Entada gigas* (reproduced with permission from Acevedo-Rodríguez 2015 onwards), (B) *Entada rheedei* subsp. *rheedei* and (C) *Entada phaseoloides* (reproduced with permission from Yang et al. 2016; arrows indicated in the original image were removed).

## References

Acevedo-Rodríguez, P. (2015, onwards). Lianas and climbing plants of the Neotropics. Retrieved from https://naturalhistory.si.edu/research/botany/research/lianas-and-climbing-plants-neotropics

Angyalossy, V., Angeles, G., Pace, M. R., Lima, A. C., Dias-Leme, C. L., Lohmann, L. G., & Madero-Vega, C. (2012). An overview of the anatomy, development and evolution of the vascular system of lianas. Plant Ecology & Diversity, 5(2), 167–182. 10.1080/17550874.2011.615574

Angyalossy, V., Pace, M. R., Evert, R. F., Marcati, C. R., Oskolski, A. A., Terrazas, T., … Baas, P. (2016). IAWA List of Microscopic Bark Features. IAWA Journal, 37(4), 517–615. 10.1163/22941932-20160151

Angyalossy, V., Pace, M. R., & Lima, A. C. (2015). Liana anatomy: A broad perspective on structural evolution of the vascular system. In S. A. Schnitzer, F. Bongers, R. J. Burnham, & F. E. Putz (Eds.), Ecology of Lianas (1st ed., pp. 251–287). Wiley. 10.1002/9781118392409.ch19

Azani, N., Babineau, M., Bailey, C. D., Banks, H., Barbosa, A. R., Pinto, R. B., … Zimmerman, E. (2017). A new subfamily classification of the Leguminosae based on a taxonomically comprehensive phylogeny: The Legume Phylogeny Working Group (LPWG). TAXON, 66(1), 44–77. 10.12705/661.3

Barbosa, A. C. F., Gerolamo, C. S., Lima, A. C., Angyalossy, V., & Pace, M. R. (2021). Polishing entire stems and roots using sandpaper under water: An alternative method for macroscopic analyses. Applications in Plant Sciences, 9(5), aps3.11421. 10.1002/aps3.11421

Caballé, C. (1986). Sur la biologie des lianes ligneuses en foret gabonaise. (Thèse de doctorat d’État en sciences). Université des Sciences et Techniques du Languedoc, Montpellier, France. Retrieved from https://agris.fao.org/search/ru/records/64735b5253aa8c8963084cf2

Carlquist, S. (1992). Anatomy of vine and liana stems: A review and synthesis. In F. E. Putz & H. A. Mooney (Eds.), The Biology of Vines (1st ed., pp. 53–72). Cambridge University Press. 10.1017/CBO9780511897658.004

Cunha-Neto, I. L. (2023). Vascular variants in seed plants—A developmental perspective. AoB PLANTS, 15(4), plad036. 10.1093/aobpla/plad036

Cunha-Neto, I. L. 2026. Plant Vascular Variants Database (v 1.0.1) [Data set]. Zenodo. 10.5281/zenodo.18704993

Cunha-Neto, I. L., Kozma, Z., Lichter-Marck, I. H., Acevedo-Rodríguez, P., & Onyenedum, J. G. (2025, preprint). Rampant convergent evolution of vascular oddities and a synnovation characterize the rapid radiation of Paullinieae lianas. bioRxiv. 10.1101/2025.07.12.664521

Cunha-Neto, I. L., & Onyenedum, J. G. (2023). Ectopic cambia: Connections between natural and experimental vascular mutants. American Journal of Botany, 110(10), e16246.10.1002/ajb2.16246

Cunha-Neto, I. L., Pace, M. R., Hernández-Gutiérrez, R., & Angyalossy, V. (2022). Linking the evolution of development of stem vascular system in Nyctaginaceae and its correlation to habit and species diversification. EvoDevo, 13(1), 4. 10.1186/s13227-021-00190-1

Cunha-Neto, I. L., S. Rossetto, E. F., Gerolamo, C. S., Hernández-Gutiérrez, R., Sukhorukov, A. P., Kushunina, M., A. Melo-de-Pinna, G. F., & Angyalossy, V. (2024). Medullary bundles in Caryophyllales: Form, function, and evolution. New Phytologist, 241(6), 2589–2605. 10.1111/nph.19342

Cunha-Neto, I. L., Snead, A. A., Landis, J. B., Callery, N. I., Husbands, A. Y., Specht, C. D., & Onyenedum, J. G. (2026). Ectopic cambia in wisteria vines are associated with the expression of conserved KNOX genes. Nature Communications, 17, 2190. 10.1038/s41467-026-68669-w

De Bary, A. H. (1877). Vergleichende Anatomie der Vegetationsorgane der Phanerogamen und Farne. Leipzig, Germany: Wilhelm Engelmann.

Dobbins, D. R., & Fisher, J. B. (1986). Wound Responses in Girdled Stems of Lianas. Botanical Gazette, 147(3), 278–289.

Fisher, J. B., & Ewers, F. W. (1989). Wound Healing in Stems of Lianas after Twisting and Girdling Injuries. Botanical Gazette, 150(3), 251–265.

Gentry, A. H. (1992). The distribution and evolution of climbing plants. In F. E. Putz & H. A. Mooney (Eds.), The Biology of Vines (1st) ed., pp. 3–50). Cambridge University Press. 10.1017/CBO9780511897658.003

Huang, X. H., Deng, T., Chen, J. T., Fu, Q. S., Zhang, X. J., Lin, N., Luo, P. R., Liu, Q., Kuai, X. Y., Peng, J. Y., Landis, J. B., Wei, Y. T., Wang, H. C., & Sun, H. (2025). Spatial distribution patterns and formation of global spermatophytes. Journal of Integrative Plant Biology, 67(10), 2668–2685. 10.1111/jipb.13923

Kaçamak, B., Réjou-Méchain, M., Grolleau, M., Loumeto, J.-J., Loubota Panzou, G. J., & Rowe, N. (2025). Linking anatomical diversity with functional life strategies in a liana community of central Africa. New Phytologist, 248(3), 1225–1244. 10.1111/nph.70413

Kraus, T., & Basconsuelo, S. (2009). Secondary root growth in Rhynchosia edulis Griseb. (Leguminosae): Origin of cambia and their products. Flora -Morphology, Distribution, Functional Ecology of Plants, 204(9), 635–643. 10.1016/j.flora.2008.09.004

Leme, C. L. D., Pace, M. R., & Angyalossy, V. (2021). The “Lianescent Vascular Syndrome” statistically supported in a comparative study of trees and lianas of Fabaceae subfamily Papilionoideae. Botanical Journal of the Linnean Society, 197(1), 25–34. 10.1093/botlinnean/boab015

Li, R., Yin, M., Yang, M., Chu, S., Han, X., Wang, M., & Peng, H. (2018). Developmental anatomy of anomalous structure and classification of commercial specifications and grades of the Astragalus membranaceus var. mongholicus. Microscopy Research and Technique, 81(10), 1165–1172. 10.1002/jemt.23111

Liu, Y., Xu, X., Dimitrov, D., Pellissier, L., Borregaard, M. K., Shrestha, N., … Wang, Z. (2023). An updated floristic map of the world. Nature Communications, 14(1), 2990. 10.1038/s41467-023-38375-y

Luna-Márquez, L., Sharber, W. V., Whitlock, B. A., & Pace, M. R. (2021). Ontogeny, anatomical structure and function of lobed stems in the evolution of the climbing growth form in Malvaceae (Byttneria Loefl.). Annals of Botany, 128(7), 859–874. 10.1093/aob/mcab105

Moya, R., Gondaliya, A. D., & Rajput, K. S. (2018). Development of successive cambia and formation of flat stems in Rhynchosia pyramidalis (Lam.) Urb. (Fabaceae). Plant Biosystems -An International Journal Dealing with All Aspects of Plant Biology, 152(5), 1031–1038. 10.1080/11263504.2017.1407376

Nair, M. N. B., & Mohan Ram, H. Y. (1990). Structure of Wood and Cambial Variant in the Stem of Dalbergia Paniculata Roxb. IAWA Journal, 11(4), 379–391. 10.1163/22941932-90000526

Nejapa, R., Cabanillas, P. A., & Pace, M. R. (2022). Cortical origin of the successive cambia in the stems of the charismatic temperate lianescent genus Wisteria (Fabaceae) and its systematic importance. Botanical Journal of the Linnean Society, 199(3), 667–677. 10.1093/botlinnean/boab091

Obaton, M. (1960). Les lianes ligneuses à structure anormale des forêts denses d’Afrique Occidentale (Thèse de doctorat). Université de Paris, Paris, France. Retrieved from https://books.google.co.id/books/about/Les_lianes_ligneuses_%C3%A0_structure_anorma.html?id=7CMBEQAAQBAJ&redir_esc=y

Pace, M. R., Lohmann, L. G., & Angyalossy, V. (2009). The rise and evolution of the cambial variant in Bignonieae (Bignoniaceae). Evolution & Development, 11(5), 465–479. 10.1111/j.1525-142X.2009.00355.x

Pagel, M. (1994). Detecting correlated evolution on phylogenies: a general method for the comparative analysis of discrete characters. Proceedings of the Royal Society of London. Series B: Biological Sciences, 255, 37–45. 10.1098/rspb.1994.0006

Qin, S., Wei, K., Cui, Z., Liang, Y., Li, M., Gu, L., … Zhang, Z. (2020). Comparative Genomics of Spatholobus suberectus and Insight Into Flavonoid Biosynthesis. Frontiers in Plant Science, 11, 528108. 10.3389/fpls.2020.528108

Quintanar-Castillo, A., Amorim, A. M., & Pace, M. R. (2025). Diversification of the stem vascular system in a clade of recent radiation and multiple habit transitions: The Bunchosia clade (Malpighiaceae). American Journal of Botany, 112(6), e70056. 10.1002/ajb2.70056

Quintanar-Castillo, A., & Pace, M. R. (2022). Phloem wedges in Malpighiaceae: Origin, structure, diversification, and systematic relevance. EvoDevo, 13(1), 11. 10.1186/s13227-022-00196-3

Rajput, K. S. (2003). Structure of Cambium and its Derivatives in the Compressed Stem of Canavalia ensiformis (L.) DC. (Fabaceae). Phyton, 43, 135–146.

Rajput, K. S. (2025). Wood anatomy and ontogeny of interxylary cambium in Canavalia cathartica Thouars, C. gladiata (Jacq.) DC. and Pueraria tuberosa (Willd.) DC. (Fabaceae). TAIWANIA, 70(2). 10.6165/tai.2025.70.230

Rajput, K. S., Moya, R., & Gondaliya, A. D. (2023). Ontogeny of multiple variants in the stems of Phaseolus lunatus L. (Fabaceae). Flora, 309, 152407. 10.1016/j.flora.2023.152407

Rami, M., Gurav, R. V., Gaikwad, S., & Rajput, K. S. (2025). Comparative study on the secondary xylem and phloem formation in three varieties of Mucuna pruriens (L.) DC (Fabaceae). Flora, 324, 152682. 10.1016/j.flora.2025.152682

Robert, E. M. R., Schmitz, N., Boeren, I., Driessens, T., Herremans, K., De Mey, J., … Koedam, N. (2011). Successive Cambia: A Developmental Oddity or an Adaptive Structure? PLoS ONE, 6(1), e16558. 10.1371/journal.pone.0016558

Rowe, N., & Speck, T. (2005). Plant growth forms: An ecological and evolutionary perspective. New Phytologist, 166(1), 61–72. 10.1111/j.1469-8137.2004.01309.x

Saddoy, J. K. C., & Cunha Neto, I. L. (2026). Fabaceae Ectopic Cambia Dataset (v.1.0.0) [Data set]. Zenodo. 10.5281/zenodo.19103807

Schenck, H. (1893). Beiträge zur Biologie der Lianen, insbesondere der in Brasilien einheimischen Arten. II. Beiträge zur Anatomie der Lianen. In A. F. W. Schimper (Ed.), Botanische Mittheilungen aus den Tropen (Vol. 5, pp. 1–271). Jena, Germany: Gustav Fischer Verlag.

Schnitzer, S. A., Kaçamak, B., Zombo, I., Pandi, V., Addo-Fordjour, P., Chen, Y. J., Filippo, A. D., Brockelman, W. Y., Nathalang, A., Gianoli, E., Garcia Leon, M. M., Bernal, B., Mackintosh, E., Mori, H., Davis, C. J., J. Peters, J. D., Zakaria, R. B., Ibarra-Manríquez, G., Sinaca-Colín, S., … T. De Deurwaerder, H. P. (2025). The Search for Champion Lianas: The Largest Lianas on Six Continents. Biotropica, 57(6), e70119. 10.1111/btp.70119

Siddiqui, S. (2025). Global patterns and drivers of species and genera richness of Fabaceae. Frontiers in Plant Science, 16, 1581814. 10.3389/fpls.2025.1581814

Solereder, H., Boodle, L. A., Fritsch, F. E., & Scott, D. H. (1908). Systematic Anatomy of the Dicotyledons: A Handbook for Laboratories of Pure and Applied Botany. Clarendon Press. Retrieved from https://books.google.com.ph/books?id=xYDwAAAAMAAJ

Sperotto, P., Roque, N., Acevedo-Rodríguez, P., & Vasconcelos, T. (2023). Climbing mechanisms and the diversification of neotropical climbing plants across time and space. New Phytologist, 240(4), 1561–1573. 10.1111/nph.19093

Struckmeyer, B. E., Binning, L. K., & Harvey, R. G. (1976). Effect of Dinitroanaline Herbicides in a Soil Medium on Snap Bean and Soybean. Weed Science, 24(4), 366–369.

Yang, S.-Z., Chen, P.-H., & Chen, J.-J. (2022). Stem cambial variants of Taiwan lianas. Botanical Studies, 63(1), 27. 10.1186/s40529-022-00358-5

Yang, S.-Z., Chen, P.-H., & Lin, K.-C. (2016). Cambial variants of liana species (Fabaceae) in Taiwan. TAIWANIA, 61(3). 10.6165/tai.2016.61.185

Zhang, J., Gao, G., Chen, J.-J., Taylor, G., Cui, K.-M., & He, X.-Q. (2011). Molecular features of secondary vascular tissue regeneration after bark girdling in Populus. New Phytologist, 192(4), 869–884. 10.1111/j.1469-8137.2011.03855.x

Zich, F. A., Hyland, B. P. M., Whiffin, T., & Kerrigan, R. A. (2020). Australian Tropical Rainforest Plants. Retrieved December 19, 2025, from https://apps.lucidcentral.org/rainforest/

