## Supplementary figures and images for "Vascular diversity in Fabaceae: evolutionary and ecological insights from a globally distributed lineage"

### Supplementary Figure 1

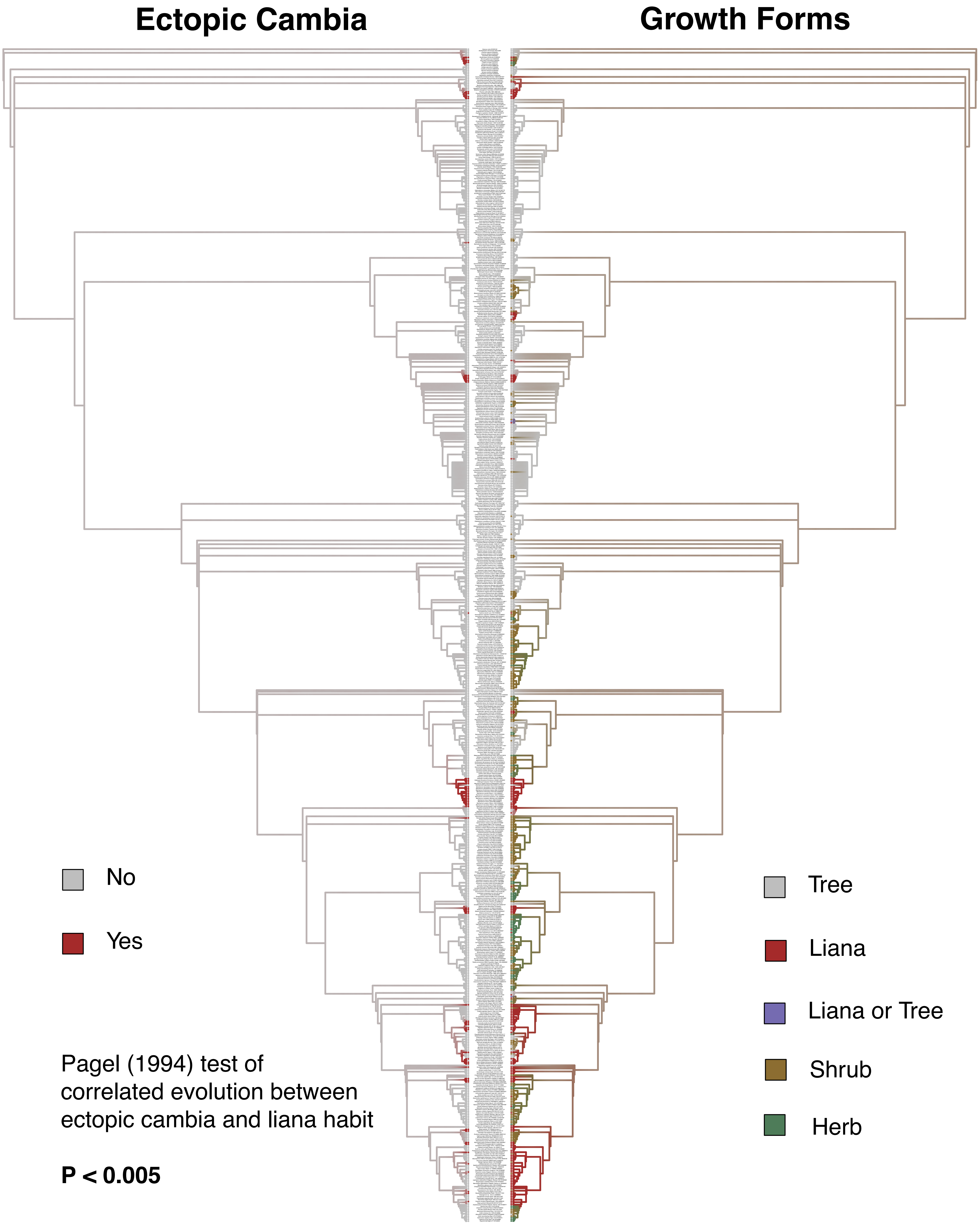

### Supplementary Figure 2

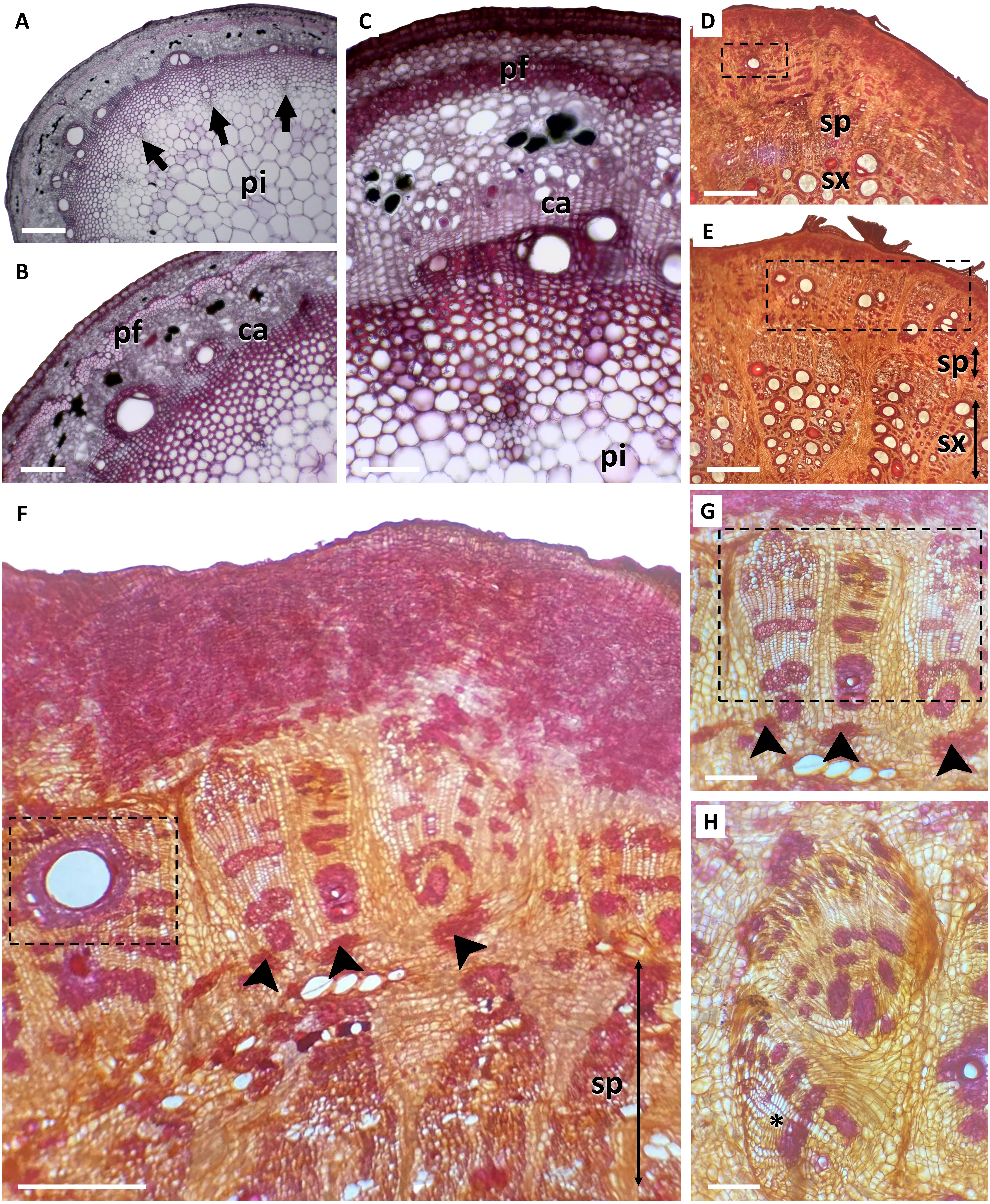

### Supplementary Figure 3

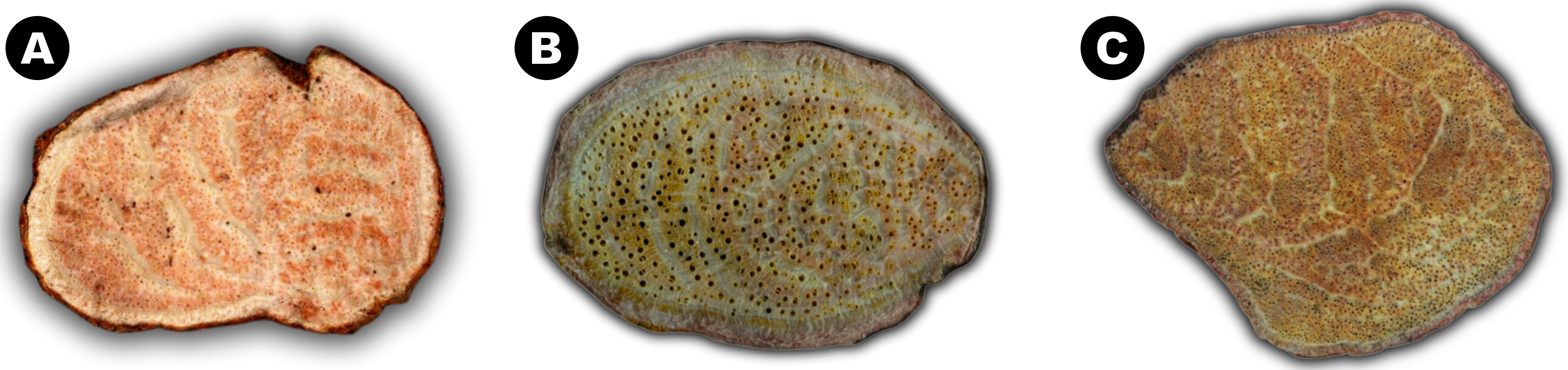


*Entada gigas* *Entada rheedei* subsp. *rheedei* *Entada phaseoloides*
